# Systematic localization of Gram-negative bacterial membrane proteins

**DOI:** 10.1101/855171

**Authors:** Anna Sueki, Frank Stein, Mikhail Savitski, Joel Selkrig, Athanasios Typas

## Abstract

The molecular architecture and function of the Gram-negative bacterial cell envelope is dictated by protein composition and localization. Proteins that localize to the inner (IM) and outer (OM) membranes of Gram-negative bacteria play critical and distinct roles in cellular physiology, however, approaches to systematically interrogate their distribution across both membranes and the soluble cell fraction are lacking. We employed multiplexed quantitative mass spectrometry to assess membrane protein localization in a proteome-wide fashion by separating IM and OM vesicles from exponentially growing *E. coli* K-12 cells on a sucrose density gradient. The migration patterns for >1600 proteins were classified in an unbiased manner, accurately recapitulating decades of knowledge in membrane protein localization in *E. coli*. For 559 proteins that are currently annotated as peripherally associated to the IM (Orfanoudaki and Economou, 2014) and display potential for dual localization to either the IM or cytoplasm, we could allocate 110 to the IM and 206 as soluble based on their fractionation patterns. In addition, we uncovered 63 cases, in which our data disagreed with current localization annotation in protein databases. For 42 of them, we were able to find supportive evidence for our localization findings in literature. We anticipate our systems-level analysis of the *E. coli* membrane proteome will serve as a useful reference dataset to query membrane protein localization, as well as provide a novel methodology to rapidly and systematically map membrane protein localization in more poorly characterized Gram-negative species.

## Introduction

The inner and outer membrane (IM and OM) of Gram-negative bacteria carry out fundamental cellular functions crucial for cell viability (Silhavy et al., 2010). The OM directly interfaces with the extracellular environment and provides a formidable physical barrier that excludes the passage of large and hydrophobic compounds (Bos et al., 2007; May and Grabowicz, 2018; Nikaido, 2003). The IM plays a vital role in ensuring selective transport of small compounds into and out of the cell, as well as sensing external cues and transducing information to adaptive transcriptional responses (Jacob-Dubuisson et al., 2018; Kuhn et al., 2017; Luirink et al., 2012). Both membranes facilitate protein translocation from the cytoplasm to the cell envelope and/or the extracellular milieu by dedicated protein machines. Proteins targeted to the IM and OM possess distinct biochemical properties and play fundamental roles in building and maintaining cell envelope integrity. This includes correct assembly of the peptidoglycan layer, which gives bacteria their cell shape and together with the OM defines their mechanical strength (Rojas et al., 2018; Typas et al., 2011). Understanding to which membrane proteins are targeted provides important insight on the physical location of their biological activity, which is particularly useful for interrogating proteins of unknown function and investigating modular envelope protein complexes (Typas and Sourjik, 2015).

Proteins localizing to the IM or OM can be sub-categorized based on their distinct biophysical properties. IM proteins typically contain α-helical transmembrane domains that mediate their integration into the phospholipid bilayer. These proteins include small molecule transporters (e.g. ABC transporters for metal ions and sugars) and large multi-protein complexes (e.g. SecYEG translocon and ATP synthase). IM proteins are diverse in their structure, function and domain localization; some contain structural domains that extend into the cytoplasmic and/or periplasmic space. In the OM, there are two distinct types of proteins: outer membrane proteins (OMPs), which are mostly composed of amphipathic β-strands that form a closed β-barrel structure, and lipoproteins, which are anchored to the OM via an N-terminal lipidated cysteine and typically contain a soluble domain that extends into the periplasmic space. Lipoproteins carry out diverse functions in the cell envelope, yet they still remain a largely poorly characterized group of proteins. Although lipoproteins have been traditionally considered to face the periplasmic space, some have been recently shown to reach the cell surface (Cowles et al., 2011; Dunstan et al., 2015; Webb et al., 2012), even by traversing through OMPs (Cho et al., 2014; Konovalova et al., 2014; Létoquart et al., 2019).

While most proteins localize to, and function at, either the IM or OM, several trans-envelope protein complexes span both membranes. Some of the larger complexes like flagella (Beeby et al., 2016) and secretion apparatuses (Deng et al., 2017) contain dedicated IM, OM and periplasmic components, whereas smaller ones such as the Tol-Pal complex (Gray et al., 2015; Lloubès et al., 2001; Petiti et al., 2019), the PBP1a/b-LpoA/B peptidoglycan synthase complexes (Egan et al., 2014; Paradis-Bleau et al., 2010; Typas et al., 2010), the efflux pump AcrAB-TolC (Du et al., 2014), the TAM translocation complex (Selkrig et al., 2012) and all TonB-dependent transport complexes (Celia et al., 2016) possess IM or OM components that can span the envelope by reaching their interaction partner in the other membrane. Moreover, in addition to integral membrane proteins, soluble proteins can associate peripherally to membranes via stable or transient protein-protein interactions with integral membrane proteins and/or with the lipid bilayer. In the case of lipoproteins, the attachment to phospholipids is covalent and part of their biosynthesis (Szewczyk and Collet, 2016). As many membrane proteins reside within protein complexes that play vital roles for cell envelope integrity, it is important to obtain a blueprint of the bacterial membrane protein composition that experimentally defines which proteins are membrane-associated and the membrane to which they are targeted. Furthermore, protein localization is closely linked to protein function, and therefore a key step in ascertaining the mode of action of membrane proteins.

Localization for most proteins in *E. coli* and other Gram-negative bacteria can be predicted quite accurately based on sequence information. Analysis of protein N-terminal signal sequences is commonly used to predict protein localization. For example, SignalP (Almagro Armenteros et al., 2019) detects signal sequences that target nascent proteins for transport across the IM via the SecYEG/SecA translocon. Similarly, TATFIND identifies substrates of the Tat translocase (Bagos et al., 2010). Interestingly, periplasmic proteins bear sufficiently distinguishable biophysical, biochemical and structural characteristics when compared to their cytoplasmic counterparts, so that one can accurately discern them with signal-sequencing agnostic machine-learning predictors (Loos et al., 2019). Moreover, several tools exist to predict membrane protein localization and topology including those that predict transmembrane α-helices and topology of IM-transmembrane proteins such as the TMHMM server (Krogh et al., 2001), as well as algorithms to predict β-barrel folding of OMPs such as PRED-TMBB (Bagos et al., 2004) and BOMP (Berven et al., 2004). *E. coli* genome databases, such as STEPdb (Orfanoudaki and Economou, 2014), Ecocyc (Keseler et al., 2017) and Uniprot (The UniProt Consortium, 2019), use information from such prediction tools, together with experimental evidence, to assign protein localization. However, the difficulties of assigning protein localization solely based on structure and signal sequence prediction can lead to mis-annotations, as it has been discussed previously (Babu et al., 2018; Orfanoudaki and Economou, 2014). Thus, as much of the current knowledge is based on prediction tools, it is of key importance to experimentally verify protein localization to clarify the predictive accuracy of the above mentioned *in silico* approaches.

Proteomic-based studies of bacterial cell membranes in the past have focused on either the IM or OM proteomes separately (Bernsel and Daley, 2009; Papanastasiou et al., 2013, 2016; Tsolis and Economou, 2017), which precludes definitive statements about the protein allocation across the two membranes and is prone to contamination from abundant soluble proteins. To systematically examine membrane protein localization in an unbiased and systematic manner, we combined sucrose gradient membrane fractionation with quantitative proteomics (Bantscheff et al., 2012). This allowed us to validate most of the predicted protein localizations, while resolving the protein localization of a large number of proteins with previously ambiguous localization, and uncovering some proteins with unexpected membrane localization, which are currently mis-annotated in databases.

## Results

### IM and OM proteome separation and quantification

To systematically assess protein localization within the bacterial cell envelope, we isolated bacterial membrane proteins by harvesting *E. coli* K-12 MG1655 grown to mid exponential phase (Figure 1A). Total membrane vesicles were purified, followed by IM and OM vesicle separation on a sucrose density gradient as previously described (Anwari et al., 2010). The sucrose gradient was then separated into eleven fractions and analyzed by immunoblot and SDS-PAGE. Effective IM and OM vesicle separation was verified using SecG and BamA antiserum (Figure 1B). Fractions 2 to 11 and the total membrane (input sample prior to fractionation) were labelled with 11-plex tandem mass tag (TMT) reagents (Werner et al., 2014) and analyzed and quantified using mass spectrometry (MS) (Figure 1A and S1A). In total, we identified 1605 common proteins across the two biological replicates, and thus proceeded to data normalization and quantification as described in the materials and methods (Figure S1A and Table S1).

**Figure 1:**
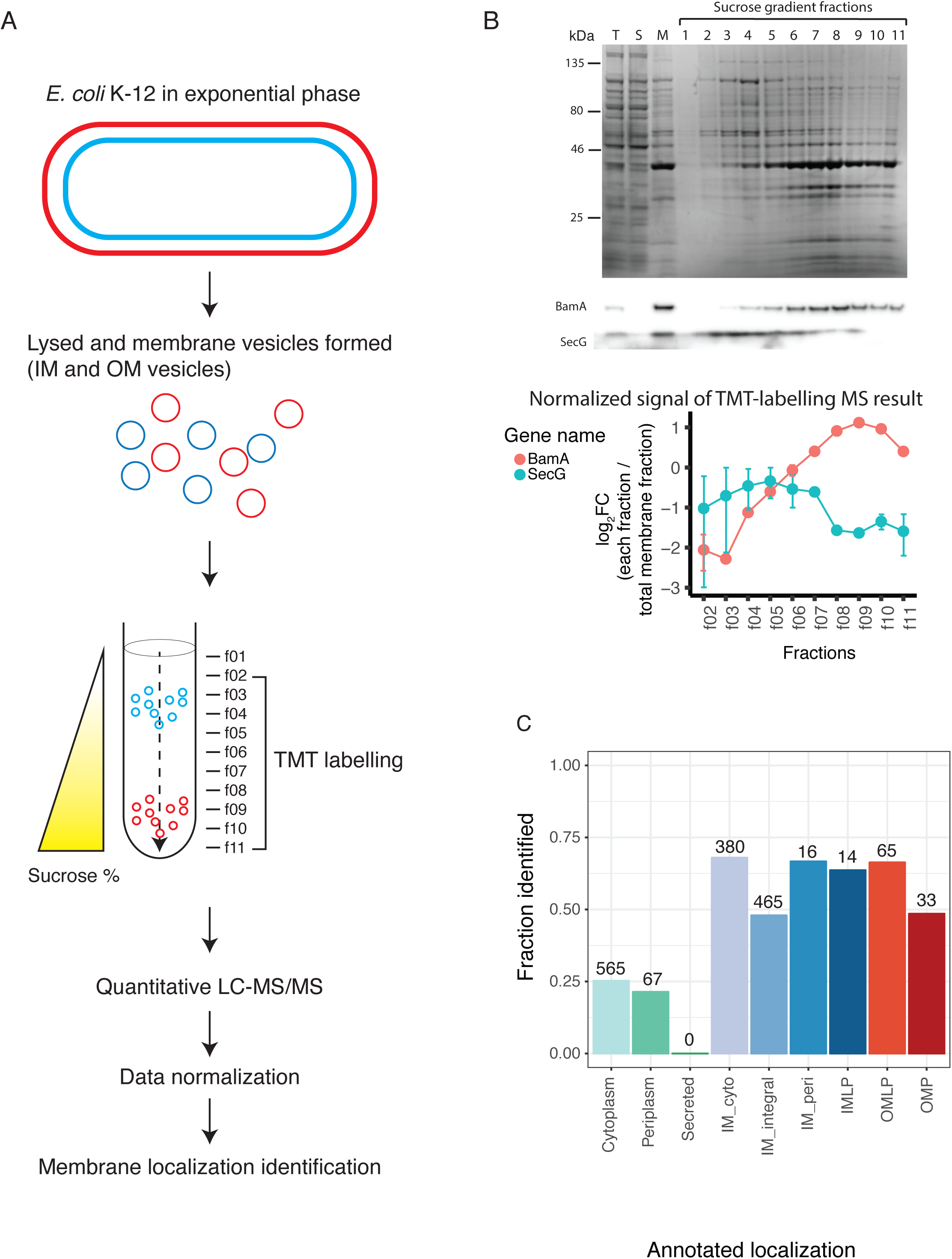
Gram-negative bacterial inner and outer-membrane fractionation quality control and membrane proteome coverage. A) Schematic illustration of the method. *E. coli* cells were harvested in exponential phase (OD_578_ ∼0.7), lysed and ultracentrifuged to collect the membrane fraction containing both OM (red) and IM (blue) vesicles. Total membrane vesicles were separated on a sucrose density gradient, which separates IM from OM vesicles based on their distinct buoyant densities. Samples were collected into 11 fractions (f01 to f11) where f02 to f11 and total membrane sample prior to fractionation were TMT-labelled and analyzed by LC-MS/MS. After data normalization (see Figure S1), membrane localization of proteins was defined. B) SDS-PAGE analysis of sucrose gradient fractionation. T: total cell lysate, S: soluble fraction upon ultracentrifugation, M: total membrane fraction prior to sucrose gradient fractionation. Upper panel: Coomassie stained gel. Middle panel: immunoblot analysis for control proteins: BamA for OM, SecG for IM (middle panel). Lower panel: TMT-labelling MS quantification result of the two control proteins (BamA and SecG). Log_2_ fold-change of each fraction / total membrane fraction for fractions (f02 to f11) for BamA and SecG. Mean with standard deviation is plotted from the two biological replicates. C) Fraction of proteins identified for each STEPdb localization category. Localization annotation is summarized in Table S2.

To assess protein abundance in each fraction, we calculated the logarithmic fold change (log_2_FC) of each sucrose gradient fraction relative to the total membrane fraction for each protein. The fractionation pattern across the 10 quantified fractions (fraction 2 to 11) was examined for each protein for each replicate (Table S1). The quantitative MS-based fractionation pattern matched immunoblot data for the control proteins, SecG for the IM and BamA for the OM (Figure 1B lower). Replicate correlation between independent experiments for all log_2_FC values was high (R = 0.77; Pearson correlation, p < 0.001, Figure S1B). Proteome coverage was analyzed by comparison to protein localization annotation from STEPdb database (Orfanoudaki and Economou, 2014) using Uniprot IDs (summarized and modified as in Table S2). For membrane protein annotation categories, we had an overall 56% coverage (973 out of 1741 proteins) with several categories reaching 70% coverage, whereas non-membrane protein categories did not exceed 25% coverage (Figure 1C). Altogether we could quantitatively assess the sucrose gradient fractionation of 1605 proteins (Table S3), covering most membrane-annotated proteins in STEPdb.

### Systematic assignment of membrane protein localization

Sucrose density gradients are conventionally analyzed by immunoblotting to compare the abundance of a given protein within a high or low sucrose density fraction (Figure 1A-B). To systematically analyze protein localization, we used the combined averages of the high and low sucrose fractions (log_2_ of f08, f09 and f10 for high, and log_2_ of f02, f03 and f04 for low) and calculated the difference between the two log_2_ averages, which we referred to as the “sucrose gradient ratio” (Table S3). High values indicate a greater abundance within higher sucrose density fractions, as expected for OM proteins. The reverse is true for IM proteins, which exhibit low values due to their enrichment within the low sucrose density fractions.

To assess whether our calculated sucrose gradient ratio reflected known protein localization, we grouped these values based on localization annotation (modified from STEPdb database, Table S2). As anticipated, most IM protein categories (i.e. IM-integral, IM-peri and IMLP) displayed a low sucrose gradient ratio, whereas the two OM protein categories, OMPs and OMLPs, showed high sucrose gradient ratios (Figure 2A). This striking concordance with curated annotations and our calculated localization confirms the accuracy of our methodology. We chose the 90th percentile of IM protein distribution (solid blue line) and the 10th percentile of OM protein distribution (solid red line) as cut-offs to define IM and OM protein localization using the sucrose gradient ratio (Figure 2B), respectively. All proteins that fell between these two cutoffs were classified as soluble proteins.

**Figure 2:**
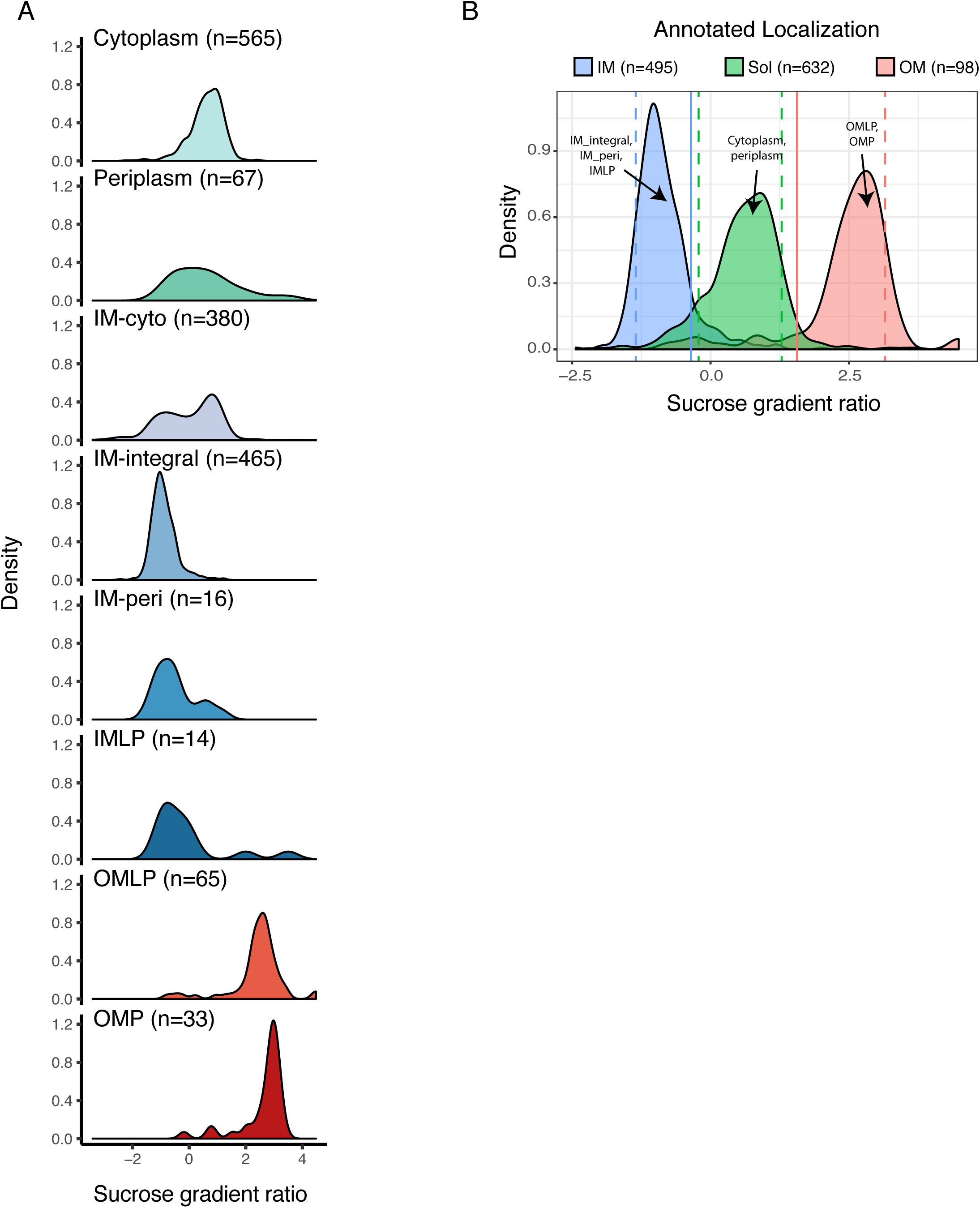
Sucrose gradient ratio separates proteins according to membrane localization. A) Smoothened distributions of sucrose gradient ratios for each protein localization category based in STEPdb (as in Figure 1C). Sucrose gradient ratio values were calculated as a difference of the average of high sucrose fractions (log_2_ of f08, f09, f10) and the average of the low sucrose fractions (log_2_ of f02, f03, f04) for each protein. B) Sucrose gradient ratio of three categories (IM, OM and Sol) grouped from A). Cytoplasmic and periplasmic proteins from (A) were grouped as soluble proteins (n=632), IM-integral and IM-peri as IM proteins (n=495) and OM and OMLPs as OM proteins (n=98). Dotted lines refer to the 10th and 90th percentiles for IM, OM and soluble proteins for each membrane localization group, respectively. The 90th percentile for IM and the 10th percentile for OM proteins are shown as solid lines as they are used as thresholds to allocate proteins to the three categories.

Interestingly, although soluble proteins were expected contaminants in our experiments, they did not always behave as expected upon sucrose density gradient fractionation. In general, soluble proteins localized in the cytoplasm and periplasm displayed midrange sucrose gradient ratios, which is in agreement with the majority of them being contaminants and non-specifically associated with either IM or OM vesicles. We noted that the IM-cyto category displayed bimodal characteristics with one peak being consistent with IM localization and another peak that aligned with soluble proteins (Figure 2A, Table S3). This suggests that the IM-cyto category of proteins referred to as “peripheral IM proteins” in STEPdb and originally described in another study (Papanastasiou et al., 2013), consists of a mixture of proteins that have clear preferential localizations either to the cytoplasm or to the IM. We therefore did not use this category for benchmarking our data, but rather kept it to later definitively allocate the primary localization of this large group of proteins. Taken together, these data show that quantitative proteomic-based analysis of sucrose gradient fractionated membrane vesicles can rapidly and systematically localize proteins to the IM or OM.

### Unbiased clustering of sucrose gradient fractionation patterns can robustly identify protein membrane localization

As the sucrose density-based fractionation patterns of IM and OM proteins were distinct (Figure 1B lower panel after normalization), we asked whether unbiased clustering of the fractionation patterns could be used to distinguish proteins according to their annotated membrane localization. Based on PCA analysis of all identified proteins, we set the number of clusters to 4 for k-means clustering analysis (Figure S2). This resulted in four groups containing the following number of proteins – cluster 1: 111, cluster 2: 650, cluster 3: 680 and cluster 4: 164 (Figure 3 left, Table S3), where each cluster exhibited distinctive fractionation patterns (Figure 3 middle). We then asked what STEPdb localization annotations are associated with proteins in the four different clusters. Proteins in cluster 1 were strongly enriched for OM proteins, cluster 2 with IM proteins, and cluster 3 and 4 with soluble proteins (Figure 3 right). Thus k-means clustering successfully grouped proteins based on their membrane localization as a function of their sucrose gradient fractionation pattern. While each cluster was dominated by a single membrane localization annotation (OM-cluster 1, IM-cluster 2 or soluble-cluster 3 and 4), some proteins were grouped into clusters that conflicted with their STEPdb annotated localization with the IM-cyto group featuring prominently in various clusters (Figure 3 right). Nevertheless, these results show that systematic and unbiased clustering of sucrose gradient fractionation patterns of the membrane proteome can be used to assign membrane localization.

**Figure 3:**
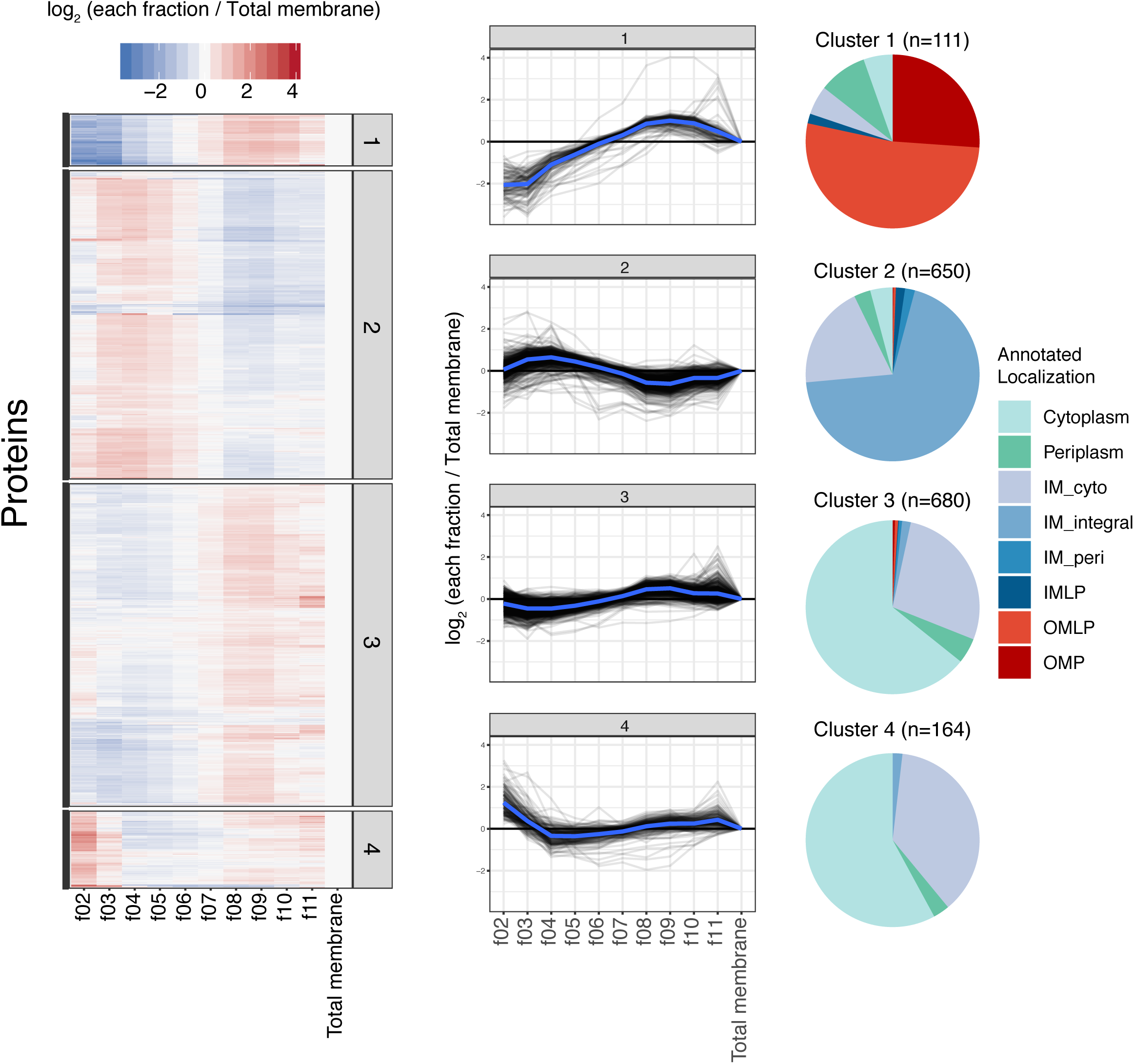
Unbiased K-means clustering of sucrose gradient fractionation patterns accurately depicts protein membrane localization. K-means clustering based on log fold-change of each fraction to total membrane sample. Left: heatmap representing each cluster patterns. Middle: fractionation pattern of all proteins in each cluster (grey) and the distribution average (blue). Right: Pie chart representing annotated localization of proteins found in each cluster. Localization annotation as in Figure 1C.

### Filtering and comparison of sucrose gradient ratio and clustering

In order to increase the confidence of our calls for protein localization, we carried out two further steps. First, we ran k-means clustering for the dataset where the two replicates were treated separately. Out of this, we identified 140 (out of 1605) proteins whose fractionation patterns between replicates resulted in clustering to different localization clusters (OM-cluster 1, IM-cluster 2 or soluble-cluster 3 and 4) for the two replicates. We reasoned that this is due to irreproducibility between the replicates and removed these proteins from further analysis. Second, we assessed the similarity of our two methods (k-means vs. sucrose gradient ratio) in assessing protein localization. To do this, we used the thresholds of sucrose gradient ratio for IM and OM defined in Figure 2B. We found a large overlap between the two methods for all three localization categories: IM, OM and soluble (Figure 4A). In total, we identified 1368 proteins (out of 1465 possible) to agree between the two methods, which we further used as our high confidence protein localization dataset. When considering all 1605 proteins, the two methods agreed in 1456 proteins (Figure S4A). In general, both quantification methods for protein localization worked well, and in combination provided more confident identification calls (true positive rates are 95% and 48% for overlap and non-overlap sets, respectively; using STEPdb annotations as true positive). The clustering method worked better than the ratio cutoffs for OM proteins, but on the other hand, cluster 4 seemed to have the most inconsistent calls for protein localization (Figure S3).

**Figure 4:**
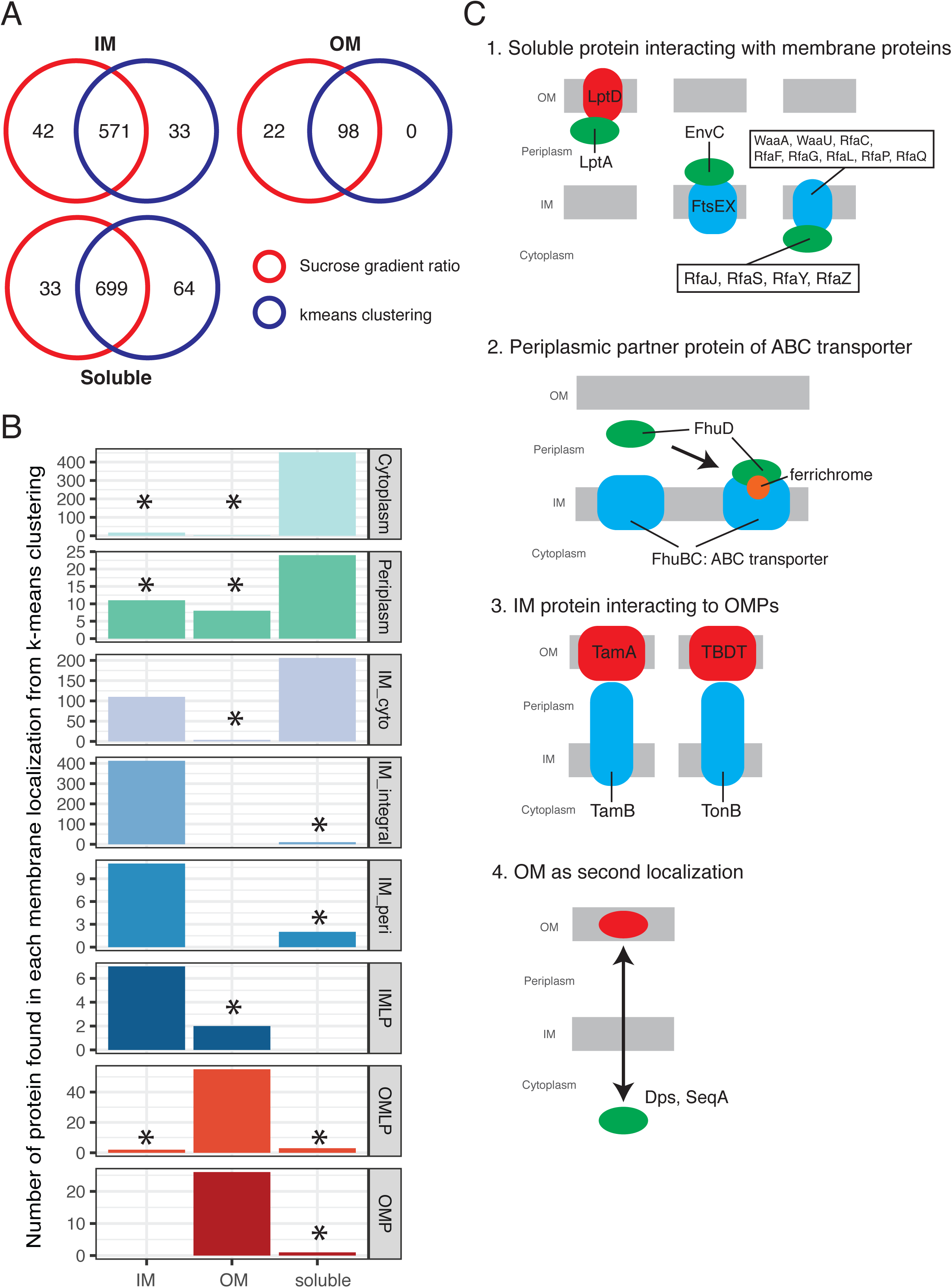
Identification of potentially mis-annotated proteins. A) Venn diagrams showing commonly identified proteins for each localization categories (IM, OM and soluble), based on our two quantification methods. Red denotes protein localizations stemming from the sucrose gradient ratios and blue refers to the k-means clustering annotation. 1465 proteins were assigned localization by the different methods. The two methods agreed in the localization of 1368 proteins, which we used as a “high-confidence” localization set. B) Comparison of identified localization with annotated localization for the 1368 proteins from panel A. Protein localizations considered not to match with STEPdb annotations are marked with *. The IM-cyto category has a bimodal distribution, likely because proteins could not be confidently allocated to either the IM or the cytoplasm in STEPdb, but can be with the experimental data provided here. C) Schematic representation explaining experimental localization of proteins not or partially matching the STEPdb annotation (besides IM-cyto): 1) Known interacting partner proteins in membranes (Left: LptA interacts with OM protein, LptD/E and fractionates as OM protein as previously reported (Chng et al., 2010), middle: EnvC interacts with IM proteins FtsEX (Yang et al., 2011), similarly for FdoG, NapG, and RseB), right: Rfa proteins in cytoplasm were identified as IM proteins, possibly due to interactions with bona-fide IM Rfa proteins). 2) Periplasmic partner of ABC transporters detected as IM proteins presumably because they bind to the IM components and transporters when transporters are active (Moussatova et al., 2008): this is for example the case for FhuD interacting with FhuBC upon substrate (ferrichrome, orange) binding. 3) Trans-envelope protein interaction is known: TamB spans the envelope and interacts with OM-located TamA (Selkrig et al., 2012), and TonB with OM-located TonB-dependent transporters, resulting in a misleading sucrose gradient fractionation (distributed equally in all fractions, thus appearing as soluble protein), which in the case of TonB has been previously shown (Higgs et al., 2002; Letain and Postle, 1997). 4) Previously shown dual localization of cytoplasmic protein in OM for Dps and SeqA (d’Alençon et al., 1999; Lacqua et al., 2006; Li et al., 2008). Full list of 63 proteins not matching with STEPdb, with 51 proteins explained can be found in Table S4.

As mentioned before, IM-cyto proteins showed a bimodal distribution of sucrose gradient ratio (Figure 2A) and could be found in multiple clusters (2, 3 and 4; Figure 3 right). Having now high-confident protein localization allocations, we could define 110 proteins as IM and 206 as soluble (Figure 4B; see also Figure S4B for all proteins, including lower confidence ones). This separation is corroborated by the melting temperatures of these two group of proteins. We have previously reported that IM proteins are more thermostable than their cytoplasmic counterparts (Mateus et al., 2018). In agreement with this, IM-cyto proteins categorized here to be soluble had similarly low melting temperatures as cytoplasmic proteins (Figure S5). In contrast IM-cyto proteins categorized as IM proteins had higher melting temperatures, albeit not as high as integral IM proteins, presumably due to their peripheral interaction rather than integral association with the IM (Figure S5). Thus, in the experimental conditions we tested, IM-cyto annotated proteins resulted in a mixture of soluble and IM proteins, which through their fractionation patterns, we could allocate their predominant protein localization.

### Identification of potentially mis-annotated proteins

STEPdb combines robust computational predictions with a wealth of experimental information to allocate protein localization in *E. coli*, and hence we used it here as a gold-standard dataset to benchmark our data and decide on thresholds for making localization calls. In doing so, we noted that a small fraction of our allocations of IM, soluble and OM proteins conflicted with their STEPdb annotations. Namely, 63 (out of 1368) high-confidence proteins, including both membrane and soluble, were found in clusters that at least partially conflicted with their corresponding STEPdb localization annotation (Table S4). We manually curated these proteins based on published literature. Firstly, we checked the localization annotation in another recent study (Babu et al., 2018). Babu *et al*. summarized different localization annotation databases and primary literature, including STEPdb, generated a score for protein localization, and then annotated protein localization accordingly in their study. Eleven proteins out of the 63 proteins we identified as mismatches agreed with the curated list in Babu *et al*. We thus conclude that the STEPdb localization for these 11 proteins was likely inaccurate (Table S4). We found corroborating evidence for 12 more such cases in literature or in other prediction databases. Importantly, in these cases rather than relying on the combined result from multiple *in silico* prediction algorithms, our data is able to provide the high-confidence experimental evidence needed to verify the localization for these proteins.

We also noted unexpected fractionation patterns for certain proteins upon sucrose density fractionation. Firstly, we detected several proteins annotated as solely periplasmic in STEPdb fractionated as membrane proteins in our experiments. Many of them have known interacting membrane partners, which is presumably the reason they co-fractionate with either the IM (EnvC, FdoG, NapG, and RseB) or the OM (LptA) (Figure 4C, Table S4). The situation was similar for periplasmic components of IM ABC transporter complexes (FhuD, PstS, and SapA) which co-fractionated with IM proteins, possibly as a consequence of a direct conditional association with IM proteins upon active transport (Moussatova et al., 2008), but were only annotated as periplasmic in STEPdb (Table S4). In total, there were 19 cases for which STEPdb had incomplete annotation.

Conversely, FecB, a known periplasmic component of an ABC transporter complex is annotated both as IM-peri (peripherally associated to IM) and periplasmic in STEPdb, but only identified as soluble in our experimental conditions. In this case, we are failing to detect the IM-association because the transporter is likely inactive in the conditions we probe, and the STEPdb annotation is more accurate (Table S4). Moreover, IM proteins known to form trans-envelope complexes (e.g. TamB and TonB) failed to cluster as either IM or OM proteins in the fractionation experiments. Overall, we could reasonably explain 51 out of the 63 cases where STEPdb and our results disagreed, out of which we could find additional information that supports our localization call (42 proteins) or the original STEPdb annotation (9 proteins) (summarized in Table S4). Overall, these findings demonstrate that our quantitative assessment of protein localization captures accurately the *in vivo* biological state.

## Discussion

We quantified the membrane proteome using TMT-labelling MS, which allowed us to experimentally identify localization in a systematic and unbiased manner for the majority of membrane proteins in *E. coli*. We verified current knowledge of membrane protein localization for proteins that was determined experimentally and/or predicted bioinformatically. The advantage of this method is that instead of assessing membrane protein localization via conventional immunoblot of sucrose density gradient fractions, quantitative proteomic approaches can be used to rapidly and quantitatively assess protein localization in an antibody-independent manner.

Comparison of our data with the curated STEPdb annotation revealed high concordance. In addition, our data provided a predominant location for a large part of the *E. coli* membrane proteome referred to as peripherally associated membrane proteins (Papanastasiou et al., 2013). STEPdb categorizes proteins that peripherally interact with the cytoplasmic face of the IM as a “peripheral IM protein”, which we referred to here for simplicity as IM-cyto (Table S2). Although we detect the majority of IM-cyto proteins (68% of 559 proteins) in the membrane fraction (which is depleted from soluble proteins), most of them are reproducibly assigned as soluble proteins according to their sucrose fractionation pattern (206 out of 316 high-confident calls). This absence of co-fractionation with the IM proteome, suggests that many of these proteins are mainly cytoplasmic in exponentially growing cells in LB, and their previous identification in membrane protein fractions in this study and others is likely because they are recurrent contaminants. We cannot exclude that some of these proteins have conditional, low affinity or transient association with the IM and proteins therein, or a small fraction of the total protein amount is at any given point associated with the IM. In contrast, about one third of the IM-cyto proteins exhibited clear IM fractionation patterns and thus can be confidently assigned as IM-associated proteins.

We found 63 proteins out of 1368 which were inconsistent with the reported localization annotation in STEPdb. We were able to explain 51 by additional literature data. Those proteins have a wrong or missing annotation in STEPdb (42) or their function/activity makes their sucrose gradient fractionation patterns misleading (9). In most cases, sucrose gradient fractionation failed to make the right call when the protein was spanning the envelope or had presumably dual membrane localization. It is likely that the new localization is also correct for most of 12 remaining proteins (Table S4). Thus, our data are helpful for improving protein localization, even for an organism as intensively studied as *E. coli*, which has been subjected to a plethora of targeted and systematic studies and researchers can benefit from carefully curated databases, such as STEPdb.

In some cases, sucrose gradient fractionation patterns were indicative of protein activity. For example, periplasmic partners of IM ABC transporters often showed sucrose gradient fractionation patterns similar to that of IM proteins (Figure 4C), which we postulate were due to their strong interaction with the cognate IM ABC transporters in the substrate-bound state (Moussatova et al., 2008). In other cases, they behaved as soluble proteins, which likely reflects the inactive state of the ABC transporter. Interestingly, trans-envelope spanning IM proteins, such as TamB and TonB, displayed fractionation patterns similar to soluble proteins. This is presumably due to strong interactions with their OM counterparts, which pull a subpopulation of the protein together with OM vesicles during ultracentrifugation. Consistent with our observations, TonB was previously found in the OM fraction upon sucrose gradient fractionation (Higgs et al., 2002; Letain and Postle, 1997). This suggests that trans-envelope IM proteins can present distinctive properties upon sucrose gradient fractionation. More broadly it implies that sucrose gradient fractionation can provide insights into the activity and mechanical strength of specific envelope complexes during different growth stages and conditions.

Not only does this work provide a resource of sucrose gradient fractionation for 1605 proteins, with 1368 proteins having their cellular localization confidently assigned, the method we present can be used for rapid and systematic characterization of membrane proteomes in different contexts – growth conditions and stages, and under different cellular perturbations. Many membrane proteins are only conditionally expressed (Mateus et al., 2018), whereas other proteins conditionally relocate in and out of membranes (Li and Young, 2012, 2015; Lim et al., 2013). Importantly, our method allows for the systematic mapping of membrane proteomes from other Gram-negative species for which protein localization annotation and transport mechanisms are less (if at all) studied.

## Supporting information

Table S1 data normalization

Table S2 protein localization

Table S3 ratio clustering

Table S4 conflicting proteins

## ACKNOWLEDGEMENTS

We thank Tassos Economou for critically reading the manuscript and providing feedback; members of Typas lab for valuable discussions; H. Tokuda for the SecG antibody; T. Lithgow for the BamA antibody; and members of the EMBL Proteomics Core Facility (PCF), especially Mandy Rettel and Dominic Helm, for assisting with sample preparation and data acquisition. We acknowledge funding from EMBL for this research. JS was supported by fellowships from the EMBL Interdisciplinary Postdoc (EIPOD) program under Marie Skłodowska-Curie Actions COFUND (grant number 291772). AS was supported by the DFG under a grant in the priority program SPP1617 and EMBL.

## DATA AVAILABILITY

The mass spectrometry proteomics data have been deposited to the ProteomeXchange Consortium via the PRIDE (Perez-Riverol et al., 2019) partner repository with the dataset identifier PXD016403.

## CODE AVAILABILITY

The code and pipelines used for data analysis are available upon request.

---

**Figure S1:**
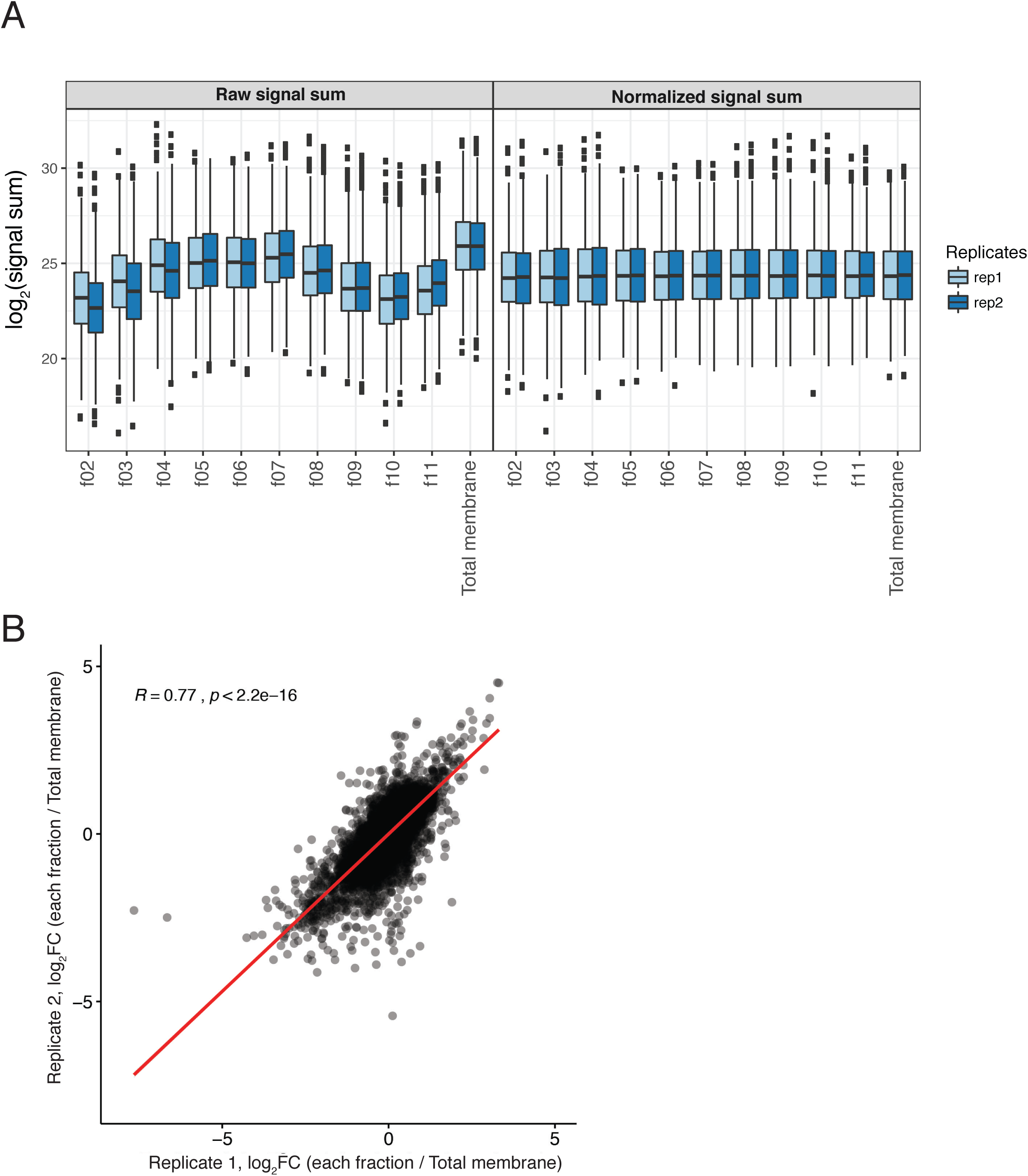
Quality control of TMT-labelling MS results. A) Boxplot representation of summed TMT reporter ion signals (signal sums) distribution across sucrose gradient fractions before and after normalization of the two biological replicates (batch-cleaned using the removeBatchEffect function from limma (Ritchie et al., 2015) and then normalized using the vsn package (Huber et al., 2002)). Box boundaries indicate the upper and lower IQR, the median is depicted by the middle boundary and whiskers represent 1.5x IQR. B) Replicate correlation of log_2_ fold-change (logFC) value of each sucrose gradient fraction to total membrane fraction (10 data points per protein, thus total 16050 data points). Pearson’s correlation shown as red line (R = 0.77, p-value < 2.2e-16).

**Figure S2:**
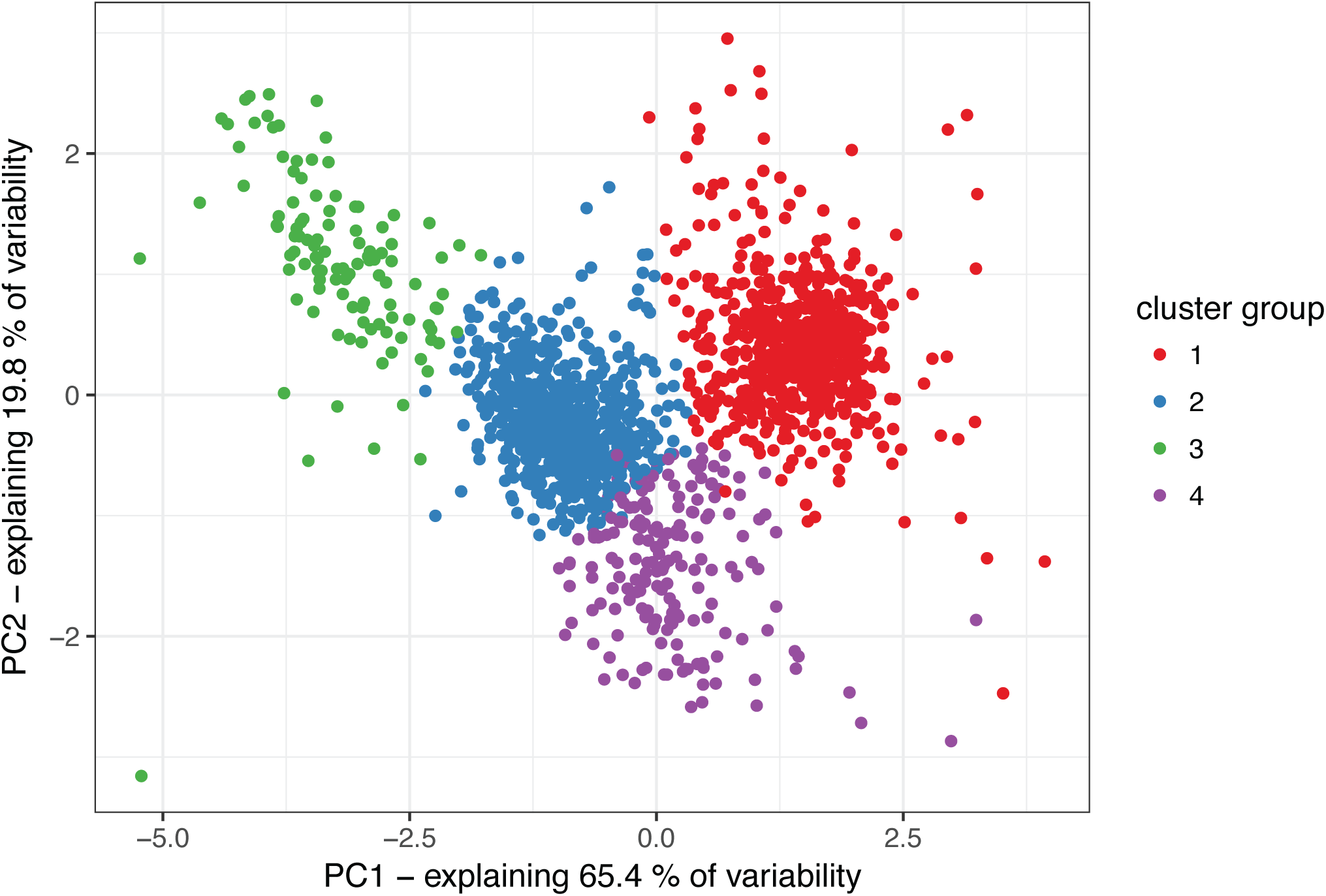
PCA analysis of fractionation patterns. Each dot represents a protein (total 1605 proteins). x-axis for PC1 (65.4% of variability), y-axis for PC2 (19.8% of variability) and color scale represents k-means clustering groups. The data used is in Table S3 (ratio of each fraction to total membrane, average of two replicates).

**Figure S3:**
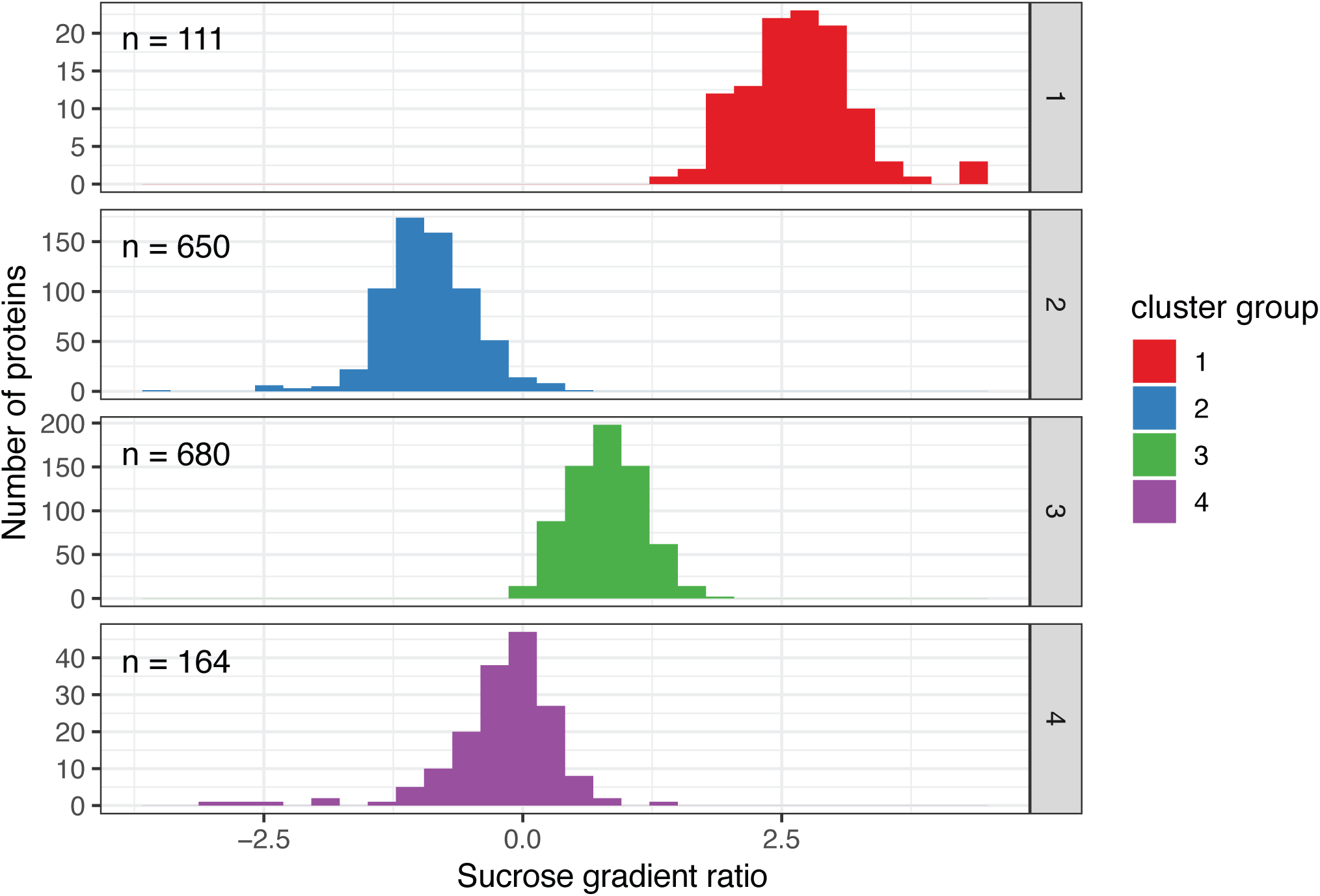
Comparison of k-means clustering to sucrose gradient ratio. Histogram of sucrose gradient ratio for each k-means cluster is shown for all proteins (1605 proteins) in the assay. Number of total protein found in each cluster are: cluster 1 – 111, cluster 2 – 650, cluster 3 – 680, cluster 4 – 164.

**Figure S4:**
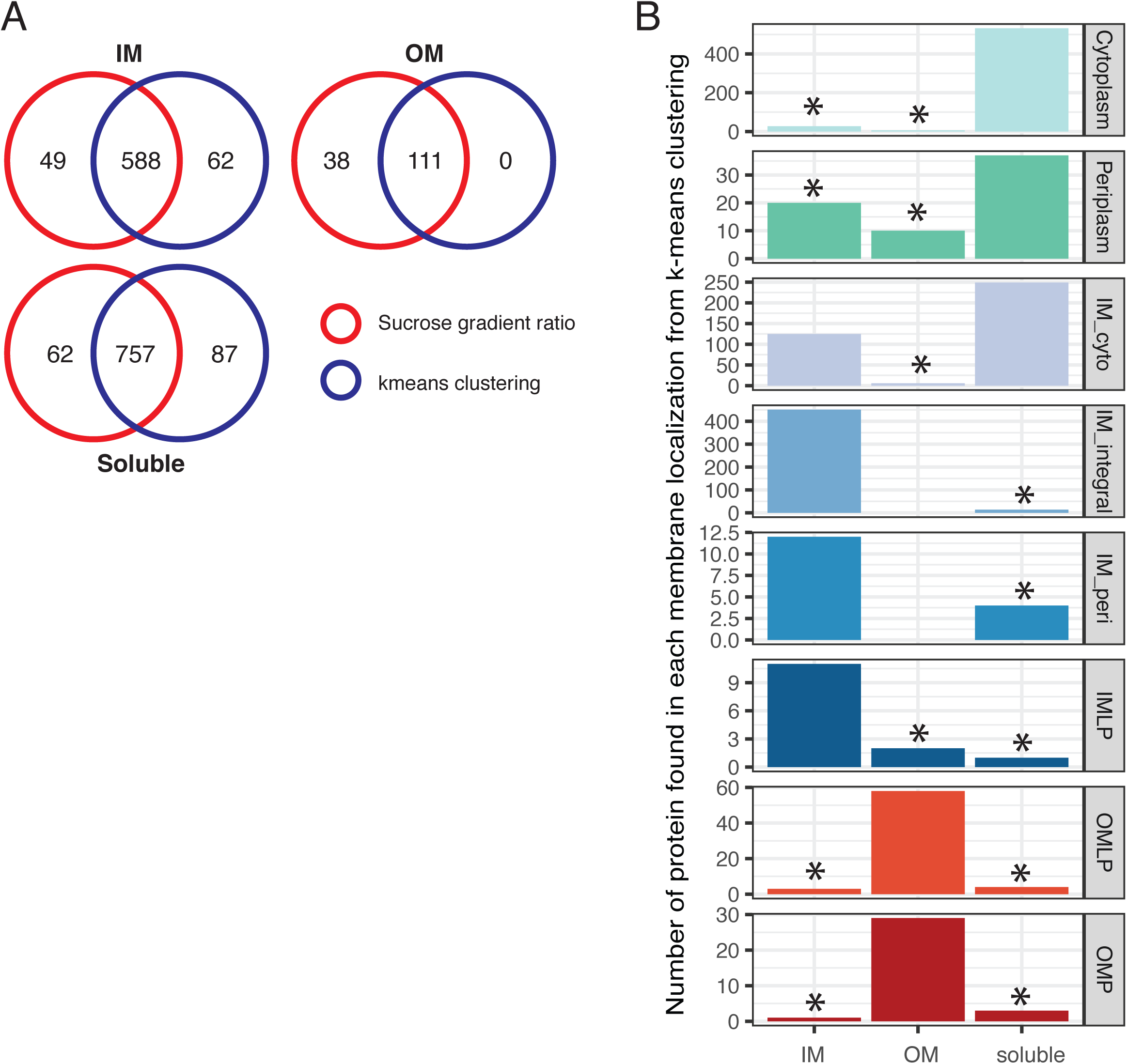
Identification of potentially mis-annotated proteins (all proteins) A) Same analysis as Figure 4A for all 1605 proteins without filtering for clustering reproducibility. Venn diagrams showing commonly identified proteins for each localization categories (IM, OM and soluble), based on our two quantification methods. Red denotes protein localizations stemming from the sucrose gradient ratios and blue refers to the k-means clustering annotation. The two methods agree in the localization of 1456 proteins. B) Same analysis as Figure 4B for all proteins without filtering for clustering reproducibility. Comparison of identified localization with annotated localization for the 1456 proteins from panel A. Protein localizations considered to be not matching with annotation are highlighted with *.

**Figure S5:**
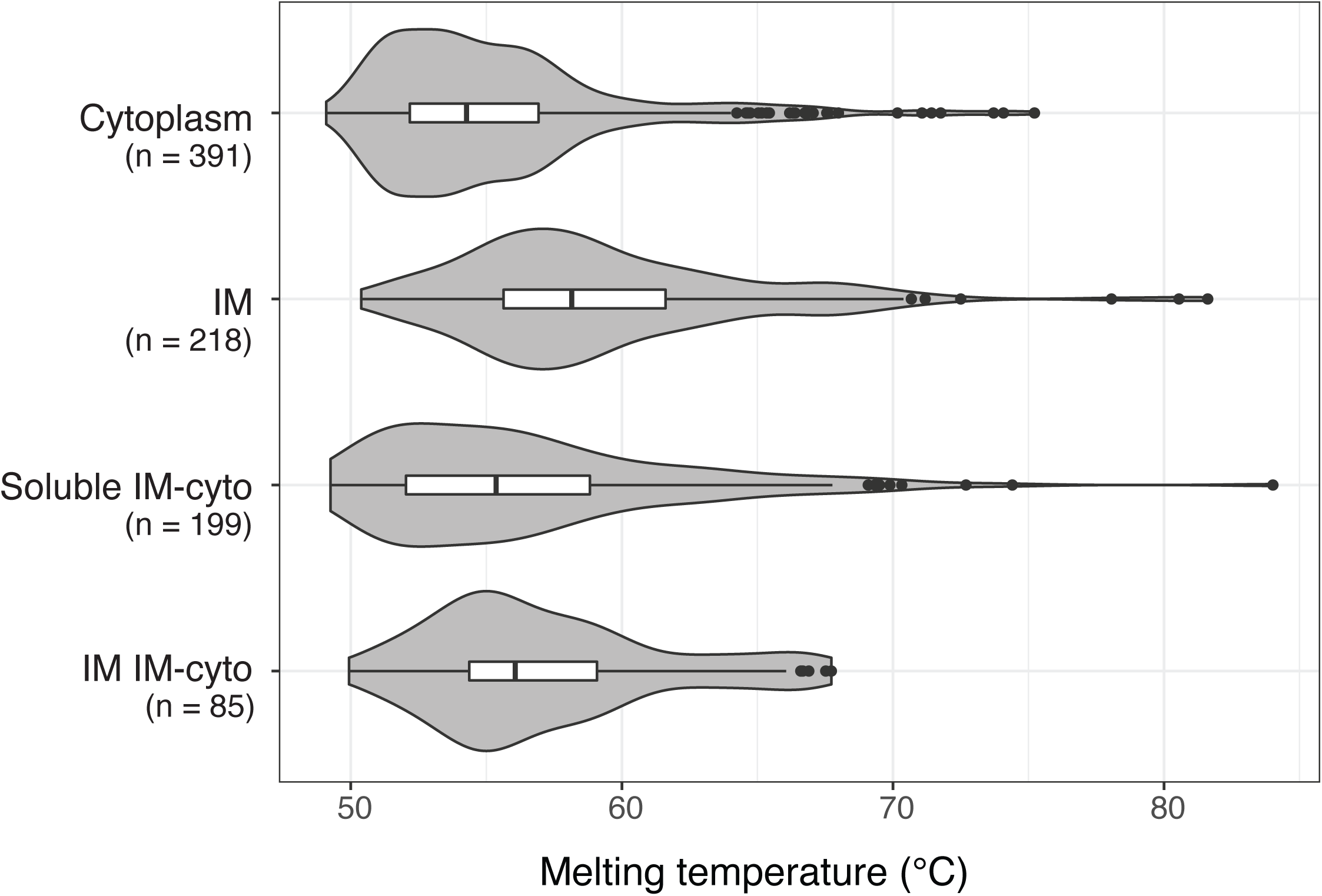
Comparison of protein localization with protein melting temperature. Violin plots encompassing box plots (plotted as in Fig. S1) of protein melting temperatures (Mateus et al., 2018) as a function of protein localization for cytoplasmic, IM and cyto-IM proteins. The latter was split in the two sub-groups we identified in our experiments. Cytoplasm (n=391), IM (n=218, IM-integral, IM-peri and IMLP combined), IM categorized IM-cyto proteins (n=85), and soluble categorized IM-cyto proteins (n=199).

## Supplemental information

Table S1. TMT-labelling MS results and normalization (signal sum values)

Table S2. Localization annotation used in this study based on STEPdb

Table S3. Membrane ratio and K-means clustering data

Table S4. Protein list non-matching STEPdb annotations & literature search summary

## Materials and Methods

### Bacterial culturing

The wild-type strain used in this study is *Escherichia coli* K-12 MG1655 Δ(*argF-lac*)U169 *rprA::lacZ* (Majdalani et al., 2002). Bacterial cells were grown for 4 generations in LB-Lennox (referred as LB herein) medium at 37°C with vigorous shaking 200 rpm, and collected for fractionation while still being in exponential growth phase, at an OD (578nm) of 0.6 – 0.8.

### Membrane vesicle isolation and sucrose density fractionation

Membrane vesicles were isolated and fractionated essentially as previously described (Anwari et al., 2010) with the following deviations. Phosphate Saline Buffer (PBS) was used as the base buffer instead of Tris. After sucrose gradient separation, 1 mL fractions were collected step-wise from the top of the gradient, yielding 11 fractionated samples that were analyzed by Coomassie staining and Western blotting using SDS-PAGE gels as described below. Fractions 2 to 11 (f02-f11), as well as an aliquot of the total input membrane sample (diluted 10 times in H_2_O), were labeled with 11-plex TMT and subjected to LC-MS/MS.

### Sample preparation and TMT labelling

Before sample preparation for MS, proteins were solubilized by adding SDS to the samples (final concentration of 1% SDS). Samples were then sonicated for 5 minutes in an ultrasonic bath, heated for 10 minutes to 80°C, and sonicated again for another 5 minutes. Disulphide bonds were reduced by incubating at 56°C for 30 minutes in 10 mM dithiothreitol (DTT) buffered with 50 mM HEPES (pH = 8.5). Reduced cysteines were alkylated by incubating 30 minutes at room temperature in dark with 20 mM 2-chloroacetamide in 50 mM HEPES buffer pH = 8.5. Samples were prepared for MS using the SP3 protocol (Hughes et al., 2014, 2019). On bead trypsin (sequencing grade, Promega, V5111) digestion was performed to an enzyme:protein ratio of 1:50 and incubated overnight at 37 °C. Digested peptides were then recovered in HEPES buffer by collecting the supernatant after magnet-based separation from the SP3 beads, and combining the second elution wash of beads with HEPES buffer. Collected peptides were labelled with TMT10plex Isobaric Label Reagent (ThermoFisher, (Werner et al., 2014)) and with 131C label (ThermoFisher) according to the manufacturer’s instructions as described below. In brief, 0.8 mg of the TMT reagents was dissolved in 42 µL of 100 % acetonitrile and 4 µL of this stock was added to the peptide sample and incubated for 1 hour at room temperature. The reaction was quenched with 5% hydroxylamine for 15 minutes at room temperature. Then the 10 samples labelled with unique TMT10plex labels were combined into one sample. The combined sample was then cleaned up using OASIS® HLB µElution Plater (Waters). The samples were separated through an offline high pH reverse phase fractionation on an Agilent 1200 Infinity high-performance liquid chromatography system which was equipped with a Germini C18 column (3 µm, 110 Å, 100 × 1.0 mm, Phenomenex). The fractionation was performed as previously described (Reichel et al., 2016). Samples were pooled in into a total of 12 fractions.

### Mass spectrometry data acquisition

Chromatography was performed using an UltiMate 3000 RSLC nano LC system (Dionex) fitted with a trapping cartridge (µ-Precolumn C18 PepMap 100, 5µm, 300 µm i.d. × 5 mm, 100 Å) and an analytical column (nanoEase™ M/Z HSS T3 column 75 µm × 250 mm C18, 1.8 µm, 100 Å, Waters). Trapping was carried out with a constant flow of solvent A (0.1% formic acid in water) at 30 µL/min onto the trapping column for 6 minutes. Subsequently, peptides were eluted via the analytical column with a constant flow of 0.3 µL/min with increasing percentage of solvent B (0.1% formic acid in acetonitrile) from 2% to 4% in 6 min, from 4% to 8% in 1 min, then 8% to 25% for a further 71 min, and finally from 25% to 40% in another 5 min. The outlet of the analytical column was coupled directly to a Fusion Lumos (Thermo) mass spectrometer using the proxeon nanoflow source in positive ion mode.

Peptides were introduced into the Fusion Lumos via a Pico-Tip Emitter 360 µm OD × 20 µm ID; 10 µm tip (New Objective) and an applied spray voltage of 2.4 kV. The capillary temperature was set at 275°C. Full mass scan was acquired with mass range 375-1500 m/z in profile mode in the orbitrap with resolution of 120000. The filling time was set at maximum of 50 ms with a limitation of 4×10^5^ ions. Data dependent acquisition (DDA) was performed with the resolution of the Orbitrap set to 30000, with a fill time of 94 ms and a limitation of 1×10^5^ ions. A normalized collision energy of 38 was applied. MS^2^ data was acquired in profile mode.

### MS data analysis

IsobarQuant (Franken et al., 2015) and Mascot (v2.2.07, (Perkins et al., 1999)) were used to process the acquired data, which was then searched against a Uniprot *Escherichia coli* proteome database (UP000000625, downloaded on 05/14/2016) containing common contaminants and reversed sequences (The UniProt Consortium, 2019). The following modifications were included into the search parameters: Carbamidomethyl (C) and TMT10 (K) (fixed modification), Acetyl (Protein N-term), Oxidation (M) and TMT10 (N-term) (variable modifications).

For the full scan (MS1) a mass error tolerance of 10 ppm, and for MS/MS (MS2) spectra of 0.02 Da was set. Further parameters were set: Trypsin as protease with an allowance of maximum two missed cleavages; a minimum peptide length of seven amino acids; at least two unique peptides were required for a protein identification. The false discovery rate on peptide and protein level was set to 0.01.

### Statistical analysis of MS data

The protein.txt output files of IsobarQuant (Franken et al., 2015) were processed using the R programming language (R Core Team, 2019). As a quality criterion, only proteins which were quantified with at least two unique peptides were used. Raw tmt reporter ion signals (signal_sum columns) were first batch-cleaned using the removeBatchEffect function from limma (Ritchie et al., 2015) and then normalized using the vsn package (Huber et al., 2002). Normalized data was clustered in 4 clusters using the kmeans function of the stat package in R.

### SDS-PAGE and Coomassie staining

Protein samples were solubilized and reduced by boiling at 95°C for 5 minutes in Laemmli loading buffer (200 mM Tris-HCl (pH=6.8), 8% SDS, 40% glycerol, 400 mM DTT, 0.02% bromophenol blue). Solubilized samples were loaded and separated in gradient gels of 4-20% acrylamide (Teo-Tricine gels from Expedeon, NXG42012) using the running buffer (Run-Blue running buffer: 0.8 M Tricine, 1.2 M Triethanolamine, 2% SDS). Bio-rad systems were used, applying 100 V per chamber. For Coomassie staining, gels were incubated in staining solution (50% methanol, 40% H_2_O, 10% acetic acid, 1 g Brilliant Blue R250 per 1 L) for 1 hour, and destained with destaining solution (40% ethanol, 10% acetic acid, 50% H_2_O) until the desirable signal was achieved. Incubations were performed at room temperature with constant moderate mixing by rocking.

### Immunoblot analysis

Proteins were separated on acrylamide gels as described above, and transferred to methanol-activated PVDF membranes (Merck, IPVH00010), using Western blot transfer buffer (3.03 g Tris, 14.4g glycine, 200 mL methanol per 1 L) for 1.5 hours at 100 V. All the incubation steps from here on were performed with constant moderate agitation by using rocking platforms. Membranes were blocked for 1 hour with 5% skim milk in TBST (20 mM Tris, 10 mM NaCl, 0.1% Tween-20), and then incubated with appropriately diluted primary antibodies (α-BamA – 1:10,000, α-SecG – 1:6,000) in 5% skim milk in TBST overnight at 4°C. After three times of 5 minutes washes with TBST, membranes were incubated for 1 hour with secondary α-rabbit antibodies conjugated with horseradish peroxidase (HRP) (GE healthcare, NA934) diluted by 1:10,000 in 5% skim milk in TBST. After these antibody incubations, membranes were washed again three times for 5 minutes with TBST. Proteins were detected by adding ECL substrate (GE Healthcare, RPN2106), then exposing and visualizing using a digital developing machine (ChemiDoc^TM^ Touch Imaging System).

### Databases

UniProt (The UniProt Consortium, 2019) was used as the source for protein ID and sequences. Protein localization information on STEPdb database (Orfanoudaki and Economou, 2014) was summarized and modified as in Table S2. Modification includes assignment of single localization for proteins with two or more localization annotations.

